# DAMM for the detection and tracking of multiple animals within complex social and environmental settings

**DOI:** 10.1101/2024.01.18.576153

**Authors:** Gaurav Kaul, Jonathan McDevitt, Justin Johnson, Ada Eban-Rothschild

## Abstract

Accurate detection and tracking of animals across diverse environments are crucial for behavioral studies in various disciplines, including neuroscience. Recently, machine learning and computer vision techniques have become integral to the neuroscientist’s toolkit, enabling high-throughput behavioral studies. Despite advancements in localizing individual animals in simple environments, the task remains challenging in complex conditions due to intra-class visual variability and environmental diversity. These limitations hinder studies in ethologically-relevant conditions, such as when animals are concealed within nests or in obscured environments. Moreover, current tools are laborious and time-consuming to employ, requiring extensive, setup-specific annotation and model training/validation procedures. To address these challenges, we introduce the ’Detect Any Mouse Model’ (DAMM), a pretrained object detector for localizing mice in complex environments, capable of robust performance with zero to minimal additional training on new experimental setups. Our approach involves collecting and annotating a diverse dataset that encompasses single and multi-housed mice in various lighting conditions, experimental setups, and occlusion levels. We utilize the Mask R-CNN architecture for instance segmentation and validate DAMM’s performance with no additional training data (zero-shot inference) and with few examples for fine-tuning (few-shot inference). DAMM excels in zero-shot inference, detecting mice, and even rats, in entirely unseen scenarios and further improves with minimal additional training. By integrating DAMM with the SORT algorithm, we demonstrate robust tracking, competitively performing with keypoint-estimation-based methods. Finally, to advance and simplify behavioral studies, we made DAMM accessible to the scientific community with a user-friendly Python API, shared model weights, and a Google Colab implementation.

**Significance:** Present deep learning tools for animal localization require extensive laborious annotation and time-consuming training for the creation of setup-specific models, slowing scientific progress. Additionally, the effectiveness of these tools in naturalistic settings is impeded by visual variability of objects and environmental diversity, hindering animal detection in complex environments. Our study presents the ’Detect Any Mouse Model’ (DAMM), a robustly validated object detector designed for localizing mice in complex environments. DAMM excels in generalization, robustly performing with zero to minimal additional training on previously unseen setups and multi-animal scenarios. Its integration with the SORT algorithm permits robust tracking, competitively performing with keypoint-estimation-based tools. These developments, along with our dissemination of DAMM, mark a significant step forward in streamlining ethologically-relevant animal behavioral studies.

## Introduction

The study of animal behavior is fundamental to disciplines such as ethology, ecology, psychology, and neuroscience. Traditional behavioral study methods, which heavily rely on manual annotation and analysis, have been pivotal in shaping our current understanding of these fields. Despite being labor-intensive and susceptible to human error, these methods have provided invaluable insights, forming the cornerstone of contemporary behavioral research.

Nonetheless, the advent of computer vision and deep learning has marked a transformative era, enabling high-throughput analyses of many processes, including animal behavior (1–3).

A crucial first step in studying animal behavior involves accurately localizing instances of animals in diverse environments. However, this task is difficult due to the inherent complexity of natural behaviors and the wide range of environments in which they are manifested (2). One major challenge for this task is intra-class visual variability, where animals within a specific category (e.g., mice), display a wide range of appearances affected by factors like posture, coat color, and social interactions. For instance, changes in posture can significantly alter the size and shape of an animal, while social interactions might result in indistinct boundaries between individuals. Additionally, environmental diversity further compounds these challenges. Varying lighting conditions and intricate structures, such as shelters and nests, can lead to occlusions, obscuring parts or the entirety of animals from view. These difficulties extend beyond animal behavior analysis and reflect core challenges in the field of computer vision.

Keypoint estimation, as implemented by tools like DeepLabCut (4) and SLEAP (5), is a prevalent approach for localizing animals in complex settings (see also (6, 7)). It involves predicting an ordered set of two-dimensional coordinates corresponding to specific body parts (e.g., left ear, snout, tail base) within an image. These sequences of keypoints form a compressed representation of animal movement, facilitating both coarse tracking and behavioral analysis. However, this approach has notable limitations. Annotating keypoints is time-consuming, labor-intensive and subjective (1, 2, 8). The subjectivity in determining the optimal placement of the keypoints can result in inconsistent and noisy predictions from the trained models, making accurate behavior prediction challenging (8). Additionally, the limited information captured by keypoint representation may not encompass all necessary details, captured by the full RGB representation, needed for accurately classifying certain behaviors (1, 8). This limitation is particularly relevant for scenarios involving subtle movements, occlusions, or interactions with other objects, such as other animals (9), nests (10), and food.

An alternative to keypoint estimation is instance segmentation. This computer vision task isolates individual objects by grouping all their pixels in an image, predicting detailed ‘masks’ that encompass the shape and location of an object. Traditionally, annotating large numbers of masks for training models has been challenging due to the required time and cost. However, recent developments in open-source interactive segmentation models have significantly eased this process. Specifically, the Segment Anything Model (SAM) has enhanced the efficiency and accuracy of mask annotation with minimal user effort (11). This advancement enables the scalable collection of instance segmentation training data, significantly expediting what is traditionally the most labor-intensive aspect of creating and validating models within the supervised learning pipeline and thereby streamlining behavioral studies.

Another major challenge in streamlining the study of animal behavior is the redundant effort and resource investments towards annotating data for training specific models that eventually do the same general task of localizing animals within complex scenes. The current strategy involves developing a unique model for each specific scenario, fitting to a specific type of animal in a specific environment. To effectively streamline this process, it is critical to develop animal detection models with strong generalization capabilities. These models should be able to generalize, or accurately make predictions on data they have not explicitly seen during training, thereby eliminating the need for additional annotation and training for end users’ experimental setups. In this work, we have adopted the standard supervised learning approach of training a deep neural network on a large and diverse dataset, a practice commonly employed to achieve robust generalization capabilities.

In this paper, we introduce the ‘Detect Any Mouse Model’ (DAMM), a robustly validated deep neural network designed for mouse detection in complex experimental setups. DAMM is engineered to generalize its detection ability to previously unseen experimental scenarios. We validate that DAMM not only excels in familiar experimental settings encountered during pretraining, but also shows exceptional generalization in entirely new scenarios without additional training, showcasing its zero-shot inference capabilities. Additionally, DAMM further improves its detection ability to new experimental setups with few training examples (≤ 50), showcasing its few-shot inference capabilities. Integrated with the SORT tracking algorithm, DAMM effectively tracks both single and multiple animals in videos, performing competitively against existing keypoint-based methods. Finally, we demonstrate DAMM’s ability to answer questions about animal behavior. DAMM’s proficiency in detection and tracking in complex multi-animal settings, combined with its generalization capabilities, represents a significant advancement in the field of animal behavior research.

## Results

### Detect Any Mouse model (DAMM) effectively localizes mice in complex environments

We aimed to develop a deep learning-based detection system capable of generalizing to novel data and accurately localizing mice in various experimental contexts with minimal additional training. Our first step was to compile a pretraining image dataset from 12,500 unique videos stored on our lab server, showcasing mice in diverse settings. This dataset, which we named the AER Lab Generalization (AER-LG) dataset (**Fig. 1A**), was diverse in terms of viewing angles (top and side), number of animals (from one to five), coat colors (black, agouti, white, and others), setup architectures (including home cages, mazes, arenas, and motor tasks), lighting conditions (varying intensities of white and red lights), and video quality (both low and high resolution, in RGB and grayscale) (**Fig. S1**).

**Fig. 1:**
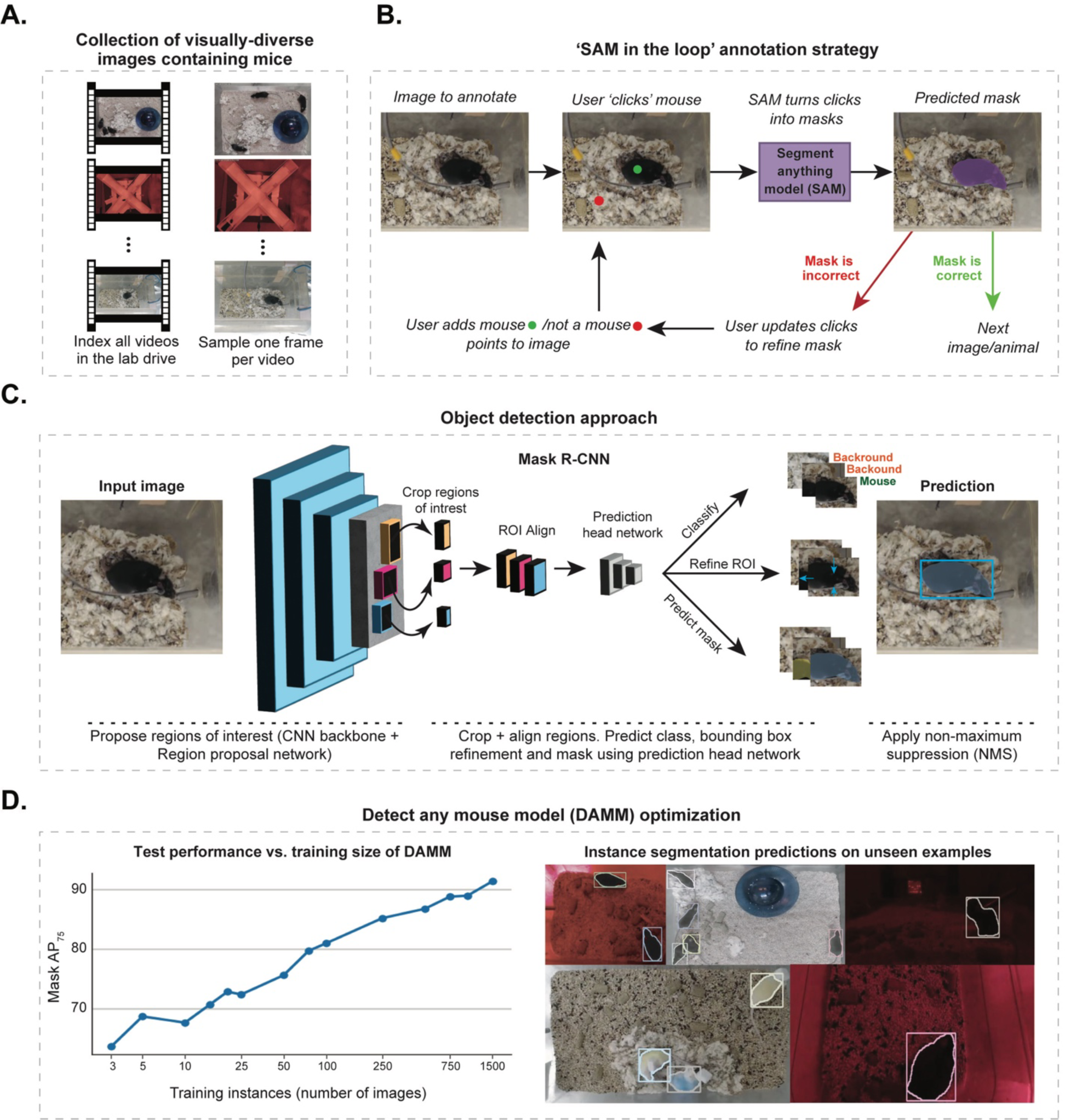
Pipeline for creating the Detect Any Mouse Model (DAMM). (A) Image dataset collection strategy. Frames were randomly extracted from an extensive archive of 12,500 videos within our laboratory (AER Lab), depicting mice in various behavioral setups. **(B)** Schematic illustration of the procedure used to generate instance segmentation masks for our pretraining dataset in a cost-effective and time-efficient manner. The schematics depicts the workflow of a graphical user interface we developed, which utilizes the Segment Anything Model (SAM) for dataset annotation. **(C)** Overview of the object detection approach, illustrating the use of Mask R-CNN, which predicts instance segments for mice within videos. **(D)** Evaluation of model performance on a test set of 500 examples. Left, COCO-style strict mask precision (IoU > .75). Right, example predictions of instance segmentation on test images. Our final pretraining dataset included 2,200 diverse images, which were utilized for training the final DAMM.

For detection and localization purposes, we trained instance segmentation models. Creating training data for this process involves annotating pixel-level delineations (masks) for each individual object instance. To streamline the labor-intensive manual annotation process, we developed a GUI integrated with SAM (11). This integration enhanced annotation efficiency by automatically generating masks using user-provided foreground/ background points (**Fig. 1B**).

To predict instance-level masks for mice in images, we trained LVIS (large vocabulary instance segmentation) (12) pretrained Mask R-CNN (13) object detectors on the AER-LG dataset annotated with our SAM GUI. Mask R-CNN is a deep neural network that performs instance segmentation in two stages. It first predicts regions of interest (ROI). Then, for all ROIs, assigns a category, decodes a mask, and refines bounding box coordinates, A process followed by non-maximum suppression to eliminate redundant instances (**Fig. 1C**). Our annotation strategy was iterative, beginning with a small training set, complemented by 200-image validation set and a 500-image test set sampled from the AER-LG dataset. By progressively increasing the training dataset with each iteration, we steadily improved our model’s test accuracy (**Fig. 1D**). After annotating 1,500 images, the model achieved a mask Average Precision at a .75 Intersection-over-Union (IoU) threshold (Mask AP75) of 92.1% on the test set (**Fig. 1D**). This high accuracy underscores the effectiveness of this supervised learning approach in training a versatile and accurate model capable of detecting mice in many complex environments. Our final model, named the Detect Any Mouse Model (DAMM), was trained on a total of 2,200 annotated images, including both validation and test sets, using the hyperparameters of our best-performing model.

### DAMM effectively detects single and multi-housed mice in complex experimental setups

We assessed the effectiveness of our DAMM in both familiar (potentially encountered during pretraining) and novel (not seen during pretraining) experimental setups, through zero-shot and few-shot evaluation procedures (**Fig. 2A**). In brief, zero-shot evaluation tests the model’s ability to detect objects in a dataset without any additional training, whereas few-shot evaluation involves fine-tuning the model with a small number (N) of annotated images from the target dataset. For these evaluations, we generated two distinct datasets: the ‘familiar’ Detect-LES dataset, compiled from five distinct setups in our lab, and the ‘novel’ Detect-PAES dataset, compiled from six publicly available datasets on the internet (and not seen during pretraining) (**Fig. 2B,C**).

**Fig. 2:**
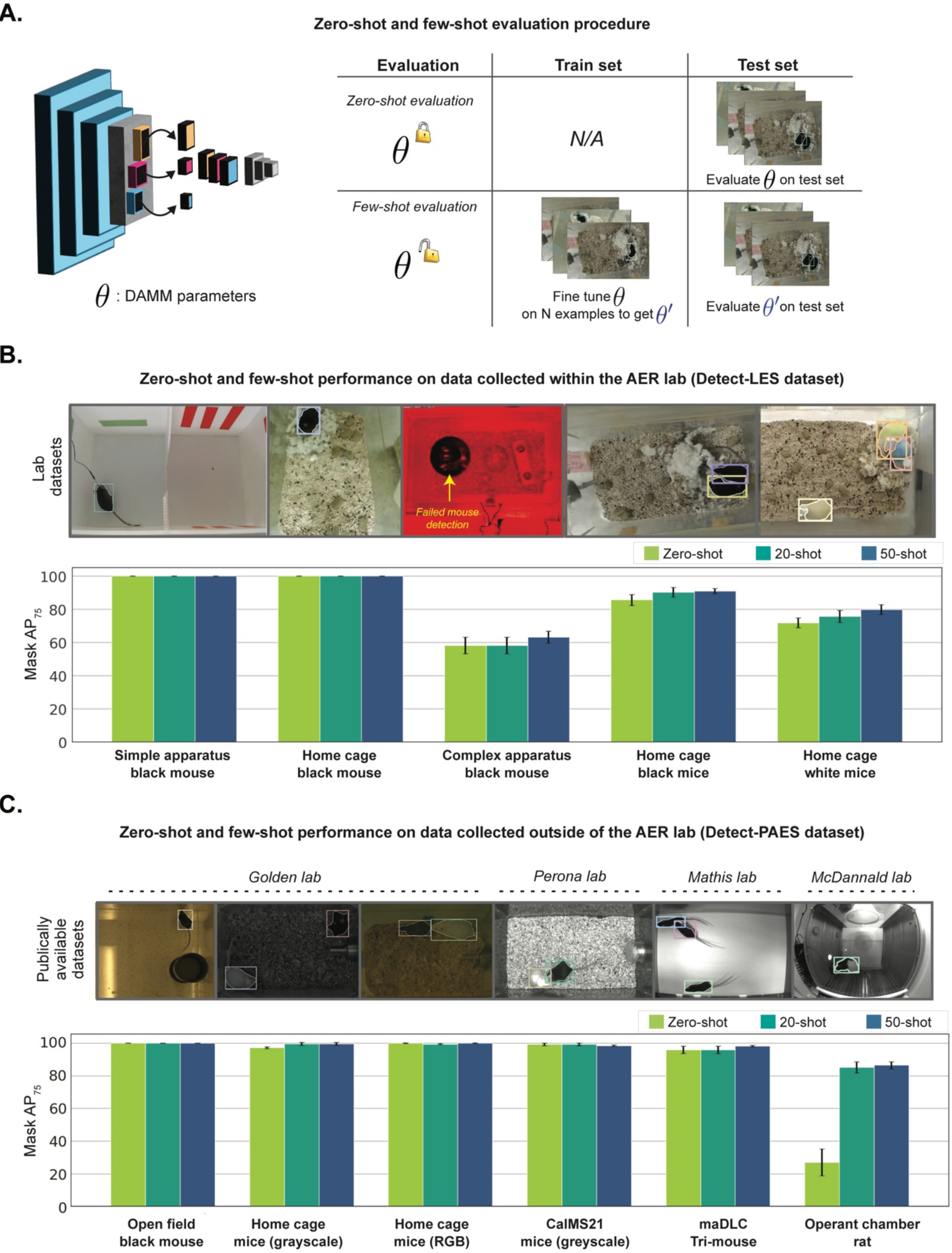
Detection performance evaluation of DAMM. (A) Schematic representation of detection evaluation procedures for two use cases: one with no further fine-tuning of model parameters (zeroshot) and another that incorporates a limited set of newly annotated examples for fine-tuning the model parameters (few-shot); θ represents model parameters. **(B)** Mask AP75 evaluation of DAMM across five unique datasets sourced from the AER Lab. The DAMM pretraining dataset may have contained frames from these five video datasets as both were sourced in-house. Each dataset contains 100 examples, with up to 50 allocated for training and 50 for testing. The mean and standard deviation of Mask AP75 are shown for each dataset across 0, 20, and 50 shot scenarios. Results are based on five randomly initialized train-test shuffles. Of note, standard deviation bars that are visually flat denote a deviation of 0. **(C)** Using the same approach as in (B), but for datasets collected outside the AER Lab. These datasets feature experimental setups that DAMM has not encountered during pretraining.

The Detect-LES dataset included three setups featuring a single black mouse: the first in a clear environment under bright light (‘Simple apparatus black mouse’), the second in a home cage with nesting material and food (‘Home cage black mouse’), and the third in an enriched environment with low red-light conditions (‘Complex apparatus black mouse’) (**Fig. 2B**).

Additionally, this dataset encompassed two setups in home cages with nesting material and food: one with two black mice (‘Home cage black mice’) and another with three differently colored mice, each equipped with head-mounted wireless devices (‘Home cage white mice’) (**Fig. 2B**). We annotated 100 randomly sampled images/frames for each setup using our SAM GUI. For each setup and few-shot scenario, we used N frames for training (with N being 0, 20, or 50), setting aside 50 images for testing.

DAMM achieved perfect accuracy without any fine-tuning (zero-shot) in both the ‘Simple apparatus black mouse’ and ‘Home cage black mouse’ setups, recording 100% Mask AP75 for zero-shot and maintaining this performance in the 20- and 50-shot scenarios (**Fig. 2B**). In the

‘Home cage black mice’ setups, the model also demonstrated high accuracy (85.6% Mask AP75) in zero-shot, with notable improvement after fine-tuning, 90.1% in the 20-shot and 90.9% in the 50-shot scenario (**Fig. 2B**). For the ‘Home cage white mice’ setup, which presented frequent occlusions, the model showed progressive improvement: starting with 71.7% Mask AP75 in zero-shot and increasing to 75.7% in the 20-shot and 79.7% in the 50-shot scenario (**Fig. 2B**). The ‘Complex apparatus black mouse’ setup posed a unique challenge due to its lighting conditions, which in some instances hindered human observers from detecting the mice as illustrated in the representative image of this setup (**Fig. 2B**). DAMM commenced with an accuracy of 58.1% Mask AP75 in zero-shot and slightly improved to 63.2% in the 50-shot scenario (**Fig. 2B**).

Collectively, these results indicate that DAMM performs exceptionally well across a variety of experimental setups relevant to laboratory mouse studies, requiring minimal training. Yet, DAMM’s performance is slightly hindered by severe occlusions and lack of visibility.

The Detect-PAES dataset, derived from publicly available videos, featured experimental setups not typical to our lab and thus not prevalent in the pretraining data, yet frequently utilized in laboratory studies (**Fig. 2C**). This strategy enabled us to assess our model’s generalization capabilities and adaptability to entirely novel scenarios. The dataset included two setups with clear backgrounds featuring either a single mouse (‘Open field black mouse’) or three mice (‘maDLC Tri-mouse’). Three other setups highlighted home cage social interactions involving black and white mice under various lighting conditions (’Home cage mice (grayscale)’, ’Home cage mice (RGB)’, and ’CaIMS21 mice (grayscale)’) (**Fig. 2C**). We also included a setup featuring a rat in an operant chamber, recorded with a fisheye lens (‘Operant chamber rat). This introduced a new rodent species and camera type not encountered during the pretraining phase (**Fig. 2C**). Overall, the Detect-PAES dataset features simpler environments compared to the Detect-LES dataset, thus posing fewer detection challenges (**Fig. 2C**). As for the Detect-LES dataset, we annotated 100 randomly sampled images/frames for each setup using our SAM GUI. For each setup and few-shot scenario, we used N frames for training (with N being 0, 20, or 50), setting aside 50 images for testing.

DAMM achieved almost perfect accuracy without any fine-tuning across the ‘Open field black mouse’, ‘Home cage mice (grayscale),’ ‘Home cage mice (RGB),’ ‘CalMS21 mice (grayscale)’ and ‘maDLC Tri-mouse’ setups, reaching 95.9-99.5% Mask AP75 in zero-shot, 20-shot, and 50-shot settings (**Fig. 2C**). In the ‘Operant chamber rat’ dataset, DAMM initially encountered challenges in the zero-shot scenario, achieving only 26.9% Mask AP75 (**Fig. 2C**). However, it showed remarkable improvement with minimal training, reaching 85.0% in 20-shot and 86.4% in 50-shot settings (**Fig. 2C**). This is particularly noteworthy since the pretrained model had never been exposed to conditions involving a fisheye lens or rats. Taken together, these findings demonstrate DAMM’s ability to generalize to novel data distributions and accurately localize rodents in various experimental contexts with minimal additional training.

### DAMM effectively detects multiple animals within challenging social and environmental conditions

We next aimed to test the extent to which our model can effectively detect mice under entirely novel, diverse, and challenging conditions, thus enabling a comprehensive evaluation of DAMM’s detection capabilities. To this end, we developed the AER Challenge dataset, which comprises 18 distinct setups each featuring three mice (**Fig. 3A-C and Fig. S2**). These setups present various challenges stemming from nontraditional recording angles, arenas cluttered with multiple objects, and low visibility. The dataset was created by combining videos from all possible combinations of three enclosure architectures (’Large cage’, ’Operant chamber’, ’Enriched cage’) (**Fig. 3A and Fig. S2**), three mouse coat colors (white, black and agouti) (**Fig. 3B and Fig. S2**) and two camera sensor qualities (entry-level and high-end, positioned at unconventional angles) (**Fig. 3C and Fig. S2**). The ’Large cage’, recorded from a top view with a slight angle, ensured consistent visibility of the mice, though occasional occlusions were caused by conspecifics. In contrast, the ’Operant chamber’, which was never encountered during pretraining, presented several challenges, including reduced visibility, increased occlusions, and object distortion all exacerbated by the camera angle and the presence of a partially observable compartment in the chamber. Lastly, the ’Enriched cage’, captured from an angled perspective, included tubes, boxes, and metal grids, which obscured the mice and allowed them to display rarely seen postures during pretraining. These scenarios introduced several computer vision challenges, including occlusions, object size variations, and within-class visual diversity. We annotated 70 randomly sampled images/frames for each setup using our SAM GUI. For each setup and few-shot scenario, we used N frames for training (with N being 0, 5, 10 or 20) and 50 frames for testing. To assess DAMM’s performance under specific experimental conditions, such as the presence of black mice, we applied marginalization, averaging results across all setups featuring the specified condition (**Fig. 3D-F**).

**Fig. 3:**
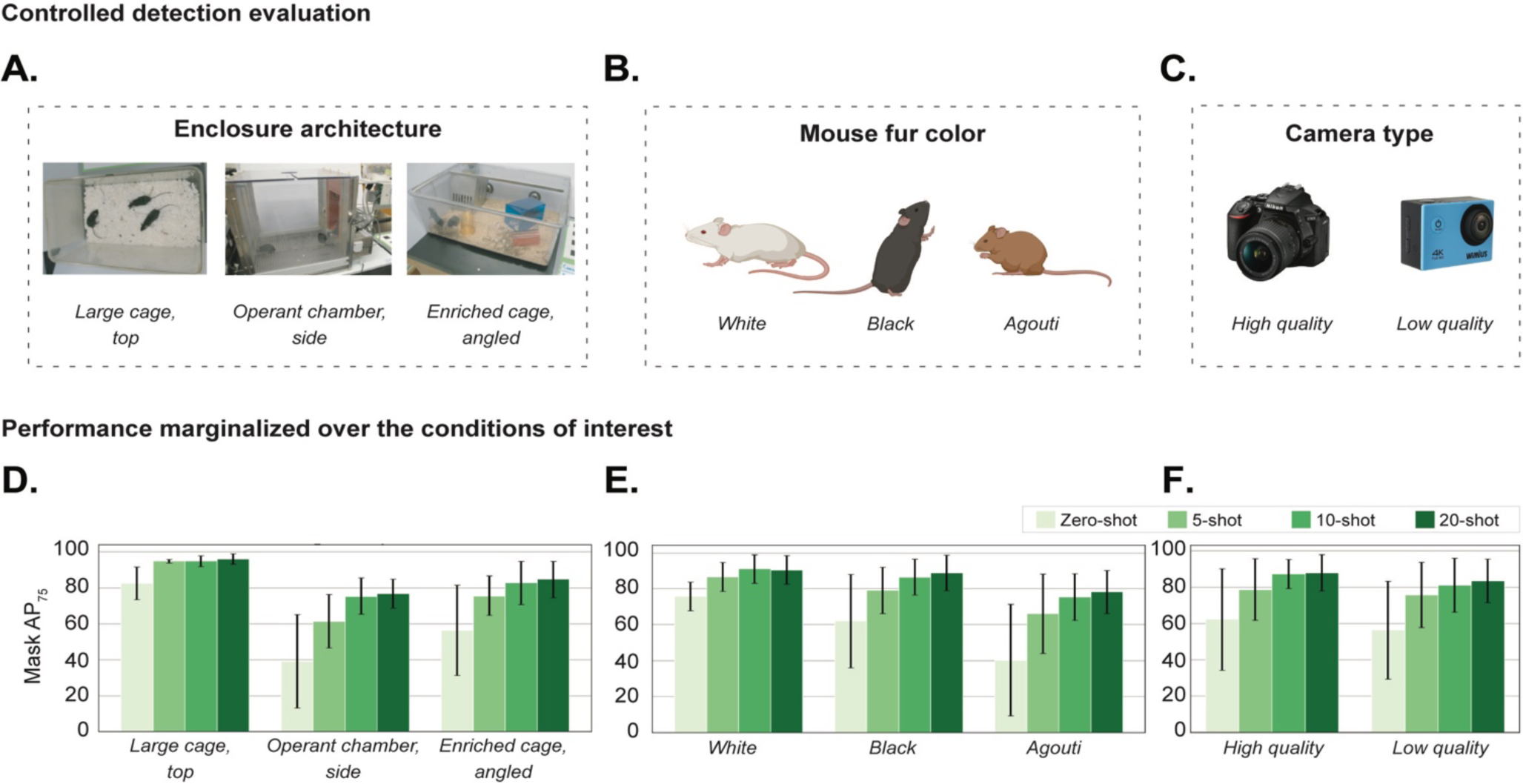
Controlled detection evaluation of DAMM. (A-C) Organization of a controlled evaluation dataset, comprising samples conditioned on three distinct groups: **(A)** environments (3 types), **(B)** mice coat colors (3 colors), **(C)** and camera types (2 types). From these categories, we generated all possible combinations, resulting in 18 mini-datasets. Each of these 18 mini-datasets contains 70 annotated frames, randomly sampled from a 5-minute video recording corresponding to the specific combination of conditions. **(D-F)** Mask AP75 average performance over all datasets containing the condition of interest, conducted for 0-shot, 5-shot, 10-shot, and 20-shot scenarios. In each scenario, we use up to 20 examples for training and 50 examples for testing.

We first examined DAMM’s capacity to detect mice under different enclosure architectures. In the ‘Large cage’ configuration, DAMM achieved high detection accuracy with a mean Mask AP75 value of 82.5% with no training and 96% with 20 additional training examples (**Fig. 3D**). In the ’Operant Chamber’ configuration, the initial zero-shot scenario accuracy was 39.2% Mask AP75, increasing dramatically to 76.8% with 20 additional training examples (**Fig. 3D**). Similarly, in the ’Enriched cage’ configuration, DAMM began with a moderate 56.4% Mask AP75 in the zero-shot scenario and improved to 84.7% with 20 additional training examples (**Fig. 3D**). Our results highlight DAMM’s exceptional performance across complex environmental conditions. Even with minimal training, DAMM consistently achieves high detection accuracy, showcasing its efficacy in the most demanding scenarios encountered in behavioral laboratory research.

We next examined DAMM’s capacity to detect mice with different coat colors (**Fig. 3B,E**). DAMM exhibited a notable ability in identifying white-coated mice, achieving a Mask AP75 of 75.7% in the zero-shot scenario and 91% in the 20-shot scenario (**Fig. 3E**). Although slightly lower than its performance with white- coated mice, DAMM also demonstrated considerable accuracy in detecting black-coated mice, starting with a mean Mask AP75 of 62% in the zero- shot scenario and steadily increasing to 88.9% in the 20-shot scenario (**Fig. 3E**). In the case of agouti-coated mice, DAMM initially showed lower accuracy, with a mean of 40.3% Mask AP75 in the zero-shot scenario. However, this significantly improved with minimal additional training, reaching 78.2% Mask AP75 in the 20-shot scenario (**Fig. 3E**). The initial low detection accuracy for aguti-coated mice is likely attributable to the underrepresentation of this coat color in DAMM’s pretraining phase. Our analysis reveals that coat color influences DAMM’s initial detection accuracy of mice, but this challenge can be effectively overcome with minimal additional annotation effort, significantly enhancing detection accuracy.

Finally, we assessed DAMM’s ability to detect mice using videos captured by both low- and high-end cameras (**Fig. 3F**). Notably, DAMM displayed a similar level of accuracy in detecting mice across different camera types (**Fig. 3F**). It started with a zero-shot performance of 56.41% Mask AP75 for low-end and 62.29% for high-end cameras, but showed a significant increase with minimal training, achieving 83.58% and 88.11% Mask AP75 respectively after annotating just 20 additional examples (**Fig. 3F**). Our findings indicate that camera quality, within the examined range, does not significantly impact DAMM’s detection accuracy, suggesting that affordable cameras are sufficient for effective detection. Collectively, our results demonstrate DAMM’s exceptional adaptability to novel, diverse, and challenging conditions with minimal annotation effort.

### DAMM effectively tracks single and multi-housed mice in complex environments with minimal training data

We proceeded to evaluate DAMM’s ability not just to detect mice in images, but also to track them in videos under diverse environmental conditions, encompassing scenarios with both single and multiple animals. To achieve this, we employed the SORT (Simple Online and Realtime Tracking) algorithm (14), which our pretrained DAMM detector integrates seamlessly with. For a comprehensive evaluation, we generated two distinct datasets: one for single-object tracking (**Fig. 4A**) and another for multi-object tracking (**Fig. 4B**). Each dataset comprised 1- minute-long video clips, with every frame and mouse annotated. For every mouse, an associated ID was annotated to track their identity throughout the video. We conducted evaluations in two scenarios: a zero-shot evaluation and a 50-shot evaluation. The evaluation metrics used were single-object tracking accuracy at an IoU threshold of 0.5 (TA50) for single-animal datasets, and multi-object tracking accuracy at IoU50 (MOTA50) for multi-animal datasets.

**Fig. 4:**
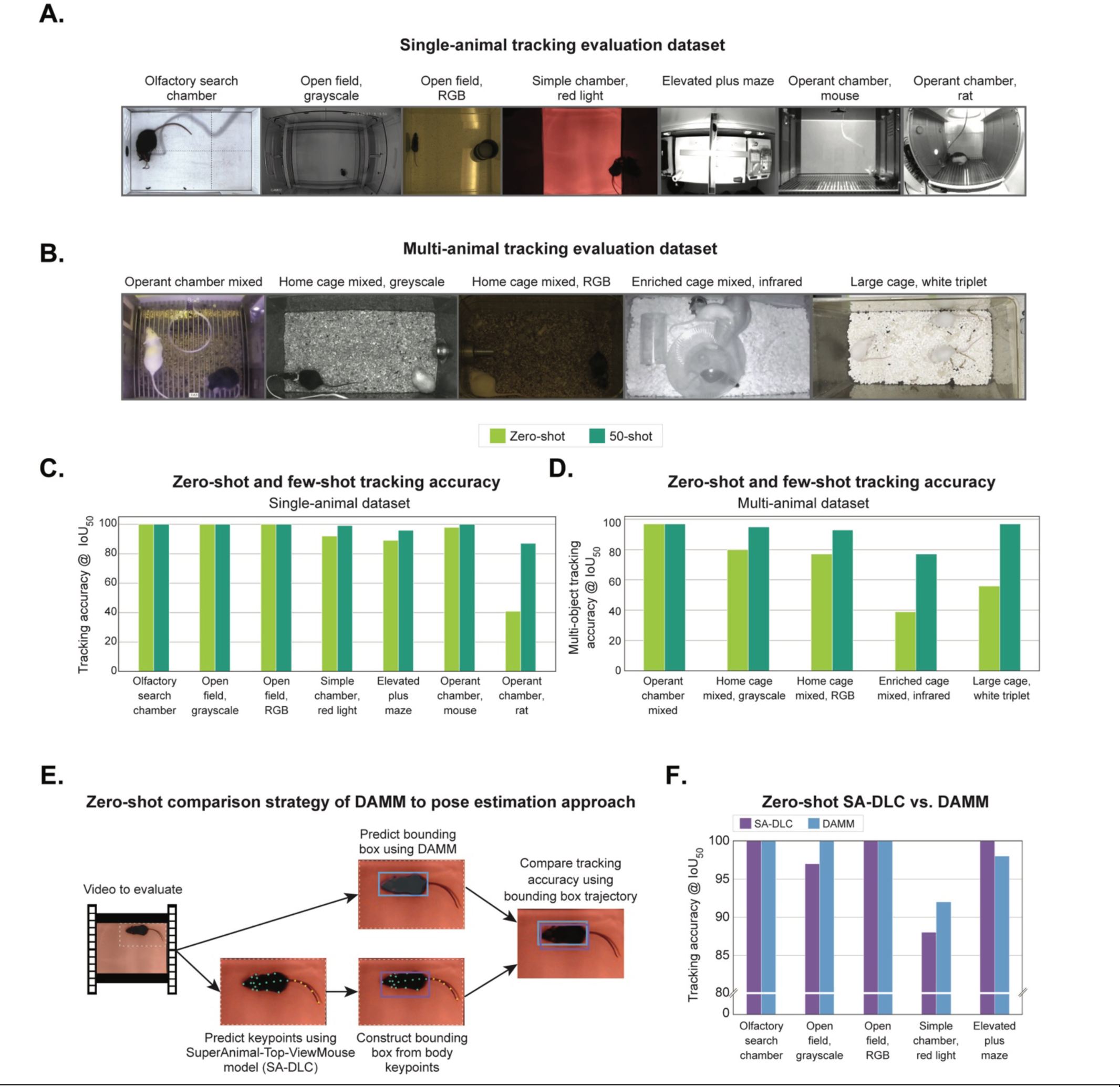
Tracking evaluation of DAMM. (**A-B)** Compilation of single-animal and multi-animal tracking evaluation datasets. Each dataset features approximately 1-minute-long videos in which the location and unique identification of every mouse are annotated throughout all frames. **(C-D)** DAMM is employed as the detection module within the Simple Online Real-time Tracking (SORT) algorithm to track mice in videos. The evaluation showcases **(C)** single-object and **(D)** multi-object tracking accuracy (IOU > .50) of DAMM for both zero-shot and 20-shot scenarios across all tracking datasets. **(E)** Comparison strategy and performance of DAMM with an existing keypoint-based-estimation mouse tracking method: the DLC SuperAnimal-TopViewMouse model. This model outputs keypoint predictions for top-view singly-housed mice. **(F)** Presented is a zero-shot tracking comparison on a subset of our previously introduced datasets which feature top-view singly-housed mice.

The single-animal tracking dataset included six experimental setups with singly-housed mice and an additional setup featuring a singly-housed rat (**Fig. 4A**). The first five mouse setups, namely ‘Olfactory search chamber,’ ‘Open field, grayscale,’ ‘Open field, RGB,’ ‘Simple chamber, red light,’ and ‘Elevated plus maze,’ all featured black coated mice and were captured from a top view. These setups varied in terms of recording distance, lighting conditions, and visual clarity. Notably, the ‘Olfactory search chamber’ included a tethered mouse, whereas the ‘Elevated plus maze’ involved a mouse navigating an elevated maze. Additionally, two operant chamber setups were recorded from a side view: one with a tethered mouse (‘Operant chamber, mouse’) and another with a rat (‘Operant chamber, rat’), the latter using a fisheye lens. All videos in this dataset were acquired through OpenBehavior Video Repository (edspace.american.edu/openbehavior/video-repository/video-repository-2/), except for the ‘Simple chamber, red light’ setup, which was recorded in the AER lab and might have been part of the pretraining data.

Overall, DAMM exhibited excellent performance in single-animal tracking across a variety of setups (**Fig. 4C**). It achieved perfect tracking accuracy, with a 100% TA50, in the ‘Olfactory search chamber,’ ‘Open field, grayscale,’ and ‘Open field, RGB’ setups without any fine-tuning, maintaining this perfect accuracy in the 50-shot scenario (**Fig. 4C**). In the ‘Simple chamber, red light,’ ‘Elevated plus maze,’ and ‘Operant chamber, mouse’ setups, DAMM showed high initial performance achieving 92%, 98%, and 89% TA50, respectively, in the zero-shot scenario (**Fig. 4C**). These accuracies further improved to 99%, 98%, and 96% TA50, respectively, after introducing 50 training examples. Notably, while DAMM’s initial accuracy for the ‘Operant chamber, rat’ setup was relatively low at 41% TA50 in the zero-shot scenario, it displayed significant improvement, reaching 87% TA50 following 50 training examples (**Fig. 4C**). These results demonstrate DAMM’s exceptional ability to generalize to novel experimental settings and rodent species and accurately track animals in videos, whether out-of-box or with minimal training effort.

The multi-animal tracking dataset encompassed five experimental setups housing multiple mice, presenting a diverse array of challenges in lighting, visibility, and animal movement (**Fig. 4B**).

This dataset included the ‘Operant chamber, mixed,’ ‘Home cage, mixed grayscale,’ ‘Home cage, mixed RGB,’ and ‘Enriched cage, mixed infrared’ setups, featuring a combination of black- and white-coated mice. The ‘Large cage, white triplet’ setup presented a unique challenge with three white mice against a white background. The ‘Enriched cage, mixed infrared’ setup, recorded in complete darkness with an infrared camera, included various enrichment items that introduced additional occlusions. The first and third videos in this dataset were sourced from OpenBehavior (edspace.american.edu/openbehavior/video-repository/video-repository-2/), the second from the Caltech Mouse Social Interactions (CalMS21) Dataset (15), and the last two from within the AER Lab.

Without training, DAMM showed variability in tracking accuracies across the different setups, yet this accuracy substantially improved with 50 training examples (**Fig. 4D**). Importantly, multi-animal tracking accuracies reflect the capacity of the model to maintain the identity of each mouse throughout the video. In the ‘Operant chamber, mixed’ setup, DAMM consistently achieved an excellent tracking accuracy with a MOTA50 of 97%, both in the zero-shot scenario and after 50 training examples (**Fig. 4D**). For the ‘Home cage, mixed grayscale’ and ‘Home cage, mixed RGB’ scenarios, initial moderate tracking accuracies in the zero-shot scenario of 77% and 80% MOTA50 rose to 93% and 95%, respectively, after 50 training examples (**Fig. 4D**). In the more challenging ‘Enriched cage, mixed infrared’ and ‘Large cage, white triplet’ setups, accuracies improved markedly from 39% and 56% MOTA50 in the zero-shot scenario to 77% and 97%, respectively, in the 50-shot scenario (**Fig. 4D**). These results underscore DAMM’s exceptional ability to not only accurately detect and track rodents but also to maintain the identities of animals within videos featuring complex multi-animal settings.

In our final evaluation, we sought to benchmark DAMM against an existing method for mouse localization, the SuperAnimal-TopViewMouse model released by DeepLabCut (SA-DLC) (16). This tool predicts body keypoint trajectories and is aimed at generalization. It is important to note that the SA-DLC model is currently released only for top-view recordings of singly-housed mice, therefore, we restricted our comparison to the subset of our datasets that match these conditions (‘Olfactory search chamber,’ ‘Open field, grayscale,’ ‘Open field, RGB,’ ‘Simple chamber, red light,’ and ‘Elevated plus maze’). To facilitate a direct comparison, we converted the predicted keypoint trajectories from the SA-DLC model into bounding box trajectories (**Fig. 4E**), allowing us to apply the same evaluation metric to both SA-DLC and DAMM (**Fig. 4F**). We found that both approaches achieved perfect tracking accuracy in the ‘Olfactory search chamber,’ and ‘Open field, RGB’ videos (**Fig. 4F**). DAMM slightly outperformed SA-DLC on two setups: in ‘Open field, grayscale’ video DAMM achieved perfect tracking accuracy with a 100% TA50, while SA-DLC scored 97% TA50, and in the ‘Simple chamber, red light,’ DAMM achieved 92% TA50, while SA-DLC scored 88% TA50 (**Fig. 4F**). SA-DLC slightly outperformed DAMM in the ‘Elevated plus maze’ video (**Fig. 4F**). SA-DLC achieved perfect tracking with a 100% TA50, while DAMM scored 98% TA50. Together, these results demonstrate DAMM’s competitive generalization capabilities over existing keypoint approaches in the top-view, single-housed animals. Moreover, DAMM extends its proficiency to include non-top view angles and complex scenarios involving multiple animals.

### DAMM can be used to answer questions about animal behavior

To showcase DAMM’s functionalities for behavioral analysis, we employed it within three experimental pipelines implemented and executed using Google Colab’s free version in the browser (**Fig. S3**). First, we utilized DAMM to process video data from a social interaction (SI) test post chronic social defeat stress (CSDS). These videos showcased black mice within clear environments (**Fig S3A and Video S1**). We employed DAMM to track the mice within the videos, utilizing zero-shot inference without the need for data annotation and training. Briefly, one day after a 10-day CSDS procedure (17), both CSDS-exposed and control mice underwent a SI test to assess the behavioral effects of chronic social defeat. Our pipeline included three main steps. Firstly, we defined zones of interest in a randomly sampled video frame. Secondly, we employed DAMM to track mice across all our videos. Finally, we computed a social interaction metric based on the proportion of time the mice’s trajectory intersected with the defined zones. We segregated the data into control and CSDS-exposed groups and replicated established distribution findings (**Fig. S3A**). Our next experimental pipeline showcases DAMM functionality in localizing mice subjected to drug treatments in obscured (under dim red light) environments (**Fig. S3B and Video S2**). In brief, this involved analyzing video data of tethered, singly-housed mice in a cluttered home-cage environment, which included nesting material and food. DAMM was employed to track the mice within the videos, utilizing few-shot inference (50 training examples). Test mice expressed the chemogenetic excitatory channel hM4Gq in a subset of lateral hypothalamic (LH) neurons, which were activated during the pre-sleep phase (10). We assessed the mice’s locomotion post-administration of either the channel ligand CNO or saline, serving as a control. DAMM’s output revealed new aspects of LH neurons’ function: besides their known role in enhancing wakefulness (10), activation of this specific subset of LH neurons also increased locomotion (**Fig. S3B**). Our third experimental pipeline showcases DAMM functionality to localize mice in multiple-animal settings using few-shot inference, fine- tunning DAMM with 100 training examples (**Fig. S3C**). Briefly, we utilized the resident-intruder assay in which an intruder mouse is inserted into the home cage of a resident mouse. DAMM robustly tracked two-colored CD-1 mice engaged in intense close-proximity interactions with frequent occlusion of one mouse by another (**Fig. S3C and Video S3**). Overall, these applications showcase our system’s functionalities in processing data across different behavioral experiments under both zero-shot and few-shot usages.

## Discussion

In this work, we release the ‘Detect Any Mouse Model’ (DAMM), a robustly validated deep neural network for detecting single and multi-housed mice within complex environments. We demonstrate DAMM’s strong generalization ability across various experimental setups and conditions, marking a significant departure from traditional methods that are reliant on labor-intensive manual annotation and the extensive training and validation procedures required to apply these models. Notably, DAMM excels in zero-shot inference, showcasing its exceptional ability to detect mice in entirely unseen experiments without any setup-specific training and achieves near-perfect performance in a variety of scenarios. When DAMM faces challenges in detecting mice out-of-box, it quickly adapts to a new experimental setup if provided with a small number of annotated training examples (≤ 50). Furthermore, we show that DAMM, when integrated with the SORT tracking algorithm, can also be used to track mice in videos across diverse environmental conditions and animal scenarios. We highlight our system’s ability to maintain the identities and locations of mice as they behave in their environment, a task that is notoriously difficult in multi-animal settings due to the risk of identity switches between closely interacting animals.

Currently, mouse detection tools that aim to generalize to new experimental setups and significantly reduce the need for experimental setup-specific fine-tuning are limited. The only publicly available mouse localization model that aims to generalize to new experimental data at the time of this work, the SuperAnimal-TopViewMouse (SA-DLC) model (16), employs a keypoint-based tracking approach that is restricted to a fixed 27-keypoint pose for singly-housed animals recorded from a top view. Not only do we demonstrate that DAMM can generalize competitively with SA-DLC for tracking mice in videos, but DAMM also extends its detection and tracking capabilities to multi-animal settings recorded from a variety of challenging scenarios— making DAMM well-suited for the full spectrum of experimental conditions neuroscientists are interested in for studying behavior. This not only underscores DAMM’s potential to significantly enhance scientific discovery but also solidifies its place in the current landscape of behavior analysis tools.

One of DAMM’s strengths lies in its accessibility. We have released our model weights and provided a Python API for our system, which is documented in a publicly available GitHub codebase (https://github.com/backprop64/DAMM), that facilitate seamless integration with existing pipelines. We also offer a set of Google Colab notebooks that showcase our system’s functionalities, as outlined in this paper. These include using DAMM zero-shot, creating and annotating a dataset, and fine-tuning DAMM. These processes are notably quick. For example, fine-tuning DAMM on free Google Colab GPUs (T4 GPU at the time of this work) only takes a few minutes. Our Colab notebooks enable users to utilize powerful behavioral analysis tools entirely in the browser, for no cost. In addition to our DAMM codebase, we have made all datasets used in this paper publicly available for further community development. In summary, DAMM is not only user-friendly but also removes the barrier of cost, enabling research labs with fewer resources to perform high-throughput behavioral analysis.

DAMM’s capabilities pave the way for several future applications. Its ability to function across diverse angles holds promise for multi-view (3D) localization applications. The speed of DAMM’s detection, which permits real-time deployment, and the on-line nature of the tracking algorithm, which only uses data from the past to make predictions about the current state of the mouse, facilitate closed-loop experiments. Additionally, the successful development process of DAMM suggests its standardized, repeatable approach could be adapted for creating generalizable detectors for other animal species.

In summary, DAMM represents a pivotal advancement in animal behavior research. Its exceptional adaptability, detection accuracy, and accessibility make it a versatile tool applicable to various experimental setups and scenarios. Moreover, its comparative performance with existing keypoint-estimation-based methods and its potential implications highlight its significance in streamlining behavioral studies and opening avenues for future research.

## Materials and Methods

### Object detection approach

We employed the Mask R-CNN architecture for instance segmentation, which detects individual objects within an image and delineates each object’s precise location with a pixel-level mask. The Mask R-CNN operates as a two-stage detector: the first stage generates predictions for regions of interest that may contain objects, while the second stage classifies these objects and refines the bounding boxes – rectangular frames outlining the exact position and size of objects – and masks associated with each object. This process involves several high-level steps: 1) Extract feature maps from the image using a convolutional neural network (CNN). 2) Predict potential object locations (regions of interest) with a Region Proposal Network (RPN) based on the feature maps. 3) Crop out regions of interest features from the predicted feature maps and resize features so they are all aligned. 4) For all ROIs, predict the object category, refine the bounding box, and generate a mask. 5) Employ a non-maximum suppression (NMS) algorithm to eliminate overlapping or low-confidence boxes.

### Tracking approach

We employed the Simple Online and Realtime Tracking (SORT) algorithm (14), specialized in single- and multi-object tracking within video streams. The SORT algorithm integrates seamlessly with our pretrained DAMM detector due to its compatibility with image-level object detectors. The algorithm involves several key steps: Initialization, Prediction, Association, and Update. In the Initialization step, objects that are repeatedly detected across frames with high overlap and are not currently being tracked are added to the set of tracked objects. The system can initiate tracking for new objects at any point during the video, provided they appear consistently across frames. In the Prediction step, the SORT algorithm estimates the next position of each tracked object based on their previous trajectories, via a Kalman filter (18). This strategy leverages kinematic information and reduces noise from the object detector. In the Association step, the SORT algorithm uses the Hungarian algorithm (19) to pair predicted locations of currently tracked objects from the Kalman filter with those provided by the object detector, optimizing matches by minimizing distance metrics such as bounding box IoU. During the Update step, the SORT algorithm refines the Kalman filter estimation for each tracked object with the matched bounding box. If there is no match, the Kalman filter independently updates using its next state prediction, effectively handling temporary occlusions. Objects are tracked until they cannot be matched to a predicted bounding box for a certain number of frames (e.g., 25). The process is iterative, cycling back to the Prediction step and continuing until the video concludes.

### Implementation details

#### Code bases

We utilized Detectron2 (20), an open-source deep learning framework developed by Meta AI, for various object detection tasks. This framework offers a user-friendly Application Programming Interface (API) for creating model architectures, managing training processes, and evaluating model performance. Additionally, for bounding box annotations in Google Colab notebooks, we used a customized version of the TensorFlow Object Detection Annotation Tool (21), adapted to fit our system’s data formats.

#### Model selection and training

To pretrain DAMM, we conducted a hyperparameter search, testing various weight decays [1e-1, 1e-2, and 1e-3] and learning rates [1e-1, 1e-2, and 1e-3]. We used the model that performed best on the validation set to evaluate/report test set performance. The final DAMM was trained using the best settings on the combined training, testing, and validation datasets. The final model was trained for 10,000 iterations using Stochastic Gradient Descent (SGD) with momentum and a batch size of 8. We started with weights from an LVIS pretrained Mask R-CNN.

For the fine-tuning of DAMM for few-shot learning on new experimental setups, we set the learning rate to 1e-1, and the weight decay at 1e-2, for 500 iterations using SGD with momentum. This fine-tuning process typically took around 5 minutes on an RTX 2080 GPU.

For the comparison to the SuperAnimal-TopViewMouse model released by DeepLabCut (16), we used predictions aggregated over scales [200,300,400,500,600] which was the only hyperparameter selected by the end-user. To construct a bounding box that is used to approximate a bounding box localization, we compute the tightest box encompassing all points, while excluding all tail points.

### Dataset Collection

#### AER Lab Generalization (AER-LG) dataset

We collected the AER-LG dataset to pretrain object detectors on diverse data encompassing a wide range of unique setups typical in behavioral studies involving mice. We compiled this dataset from a lab drive containing a rich repository of about 12,500 behavioral experiment videos collected over seven years. For each video, we randomly sampled one frame, and after a curation process, we selected 10,000 diverse images for annotation.

During the annotation phase, we employed an iterative process, initially annotating a small training set alongside a 200-image validation set and a 500-image test set. With each iteration, we expanded the training set as we annotated additional batches of images from the remaining unannotated images. After annotating each batch, we trained our object detectors and assessed their performance on the test set. This cycle of annotating and training continued, with successive additions to the training set, until performance converged. Ultimately, we annotated 1,500 images, reaching an accuracy of 92% mask Average Precision (AP) at a 75% IoU threshold on our test set. Our final dataset contains 2,200 images (**Fig. S1**), all annotated using the SAM annotation tool (see below).

#### Lab Experimental Setups (Detect-LES) dataset

To evaluate the DAMM detector in experimental setups typical of our lab, we collected the Detect-LES dataset. The original videos in this dataset might have been previously encountered by the DAMM detector during its pretraining phase. To facilitate a thorough evaluation, we constructed a series of five mini-datasets, each corresponding to videos originating from different downstream experimental setups stored on our lab server. The first mini-dataset featured a single mouse in a simple, brightly lit environment. The second, third, and fourth mini-datasets depicted a single black mouse, two black mice, and three colored mice, respectively, in a home cage under white light containing bedding, nesting material, and food. The fifth mini-dataset featured a single black mouse in a large enclosure, which included various enrichment objects such as a running wheel, and was recorded under dim red light. From these videos, we randomly sampled 100 frames. These frames were then annotated using our SAM GUI (see below).

#### Publicly Available Experimental Setups (Detect-PAES) dataset

To evaluate the DAMM detector on setups not encountered during its pretraining, we collected the Detect-PAES dataset using publicly available video data. The collection process mirrored that of the Detect-AER, with the key difference being the source of videos–internet sources instead of our lab. We acquired a total of six videos. Three videos were donated by Sam Golden and acquired from the OpenBehavior Video Repository (edspace.american.edu/openbehavior/video-repository/video- repository-2/): one depicting a single mouse in an open field (‘Open field black mouse’), and two showcasing a home cage social interaction setup with a black and white mouse, one recorded in greyscale (’Home cage mice (grayscale)’), and the other in RGB (’Home cage mice (RGB)’).

Additionally, we selected a video from the CalMS21 dataset (15), featuring a home cage social interaction setup with a black and white mouse, recorded in greyscale (’CaIMS21 mice (grayscale)’). From the maDLC Tri-mouse dataset (22), we curated a mini-dataset, which uniquely provided images rather than videos, allowing us to directly sample 100 random images. Finally, we included a setup donated by Michael McDannald, featuring a rat in an operant chamber recorded with a fisheye lens, also available through OpenBehavior Video Repository. We randomly sampled 100 frames for each mini-dataset. These frames were subsequently annotated using our SAM annotator (see below).

#### AER Challenge dataset

To assess the performance of DAMM under controlled conditions, with a focus on variation in image resolution, mouse coat color, and enclosure architecture, we created the AER Challenge dataset. This dataset consists of videos that were created post DAMM pretraining, utilized arenas not previously used for pretraining and were taken from non-standard angles (see below), ensuring their novelty to our system. We organized the dataset around three key variables: image quality (Entry-level, that costs in the order of tens of dollars: Explore One Action camera. High-end, that costs in the order of hundreds of dollars: Nikon d3500 DSLR camera), mouse coat color (white, black, and aguti), and enclosure architecture. The enclosures included a ‘Large cage’ with bedding (34 cm x 24 cm x 20 cm), an ‘Operant chamber’ with a metal grid floor and red walls (30 cm x 32 cm x 29 cm), and an ‘Enriched cage’ with bedding and toys (40 cm x 30 cm x 20 cm). Our objective in recording video data from non-standard angles was to assess the effectiveness of our system in tracking mice across diverse viewpoints, addressing key challenges in computer vision, such as occlusions, variations in object size, and within-class visual variability. We filmed a 5-minute video for each of the 18 possible combinations (2 x 3 x 3) of these variables. From each video, we randomly sampled 70 frames, which we annotated using our SAM annotator tool (see below).

#### Single- and Multi-Animal Tracking datasets

To evaluate DAMM’s ability to track mice within videos, we compiled two tracking datasets. Unlike our detection datasets, which are composed of annotated images, our tracking datasets consist of annotated videos. In these datasets, each data point is a video with every frame and mouse annotated. Additionally, for every mouse, an associated ID is used to maintain the object’s identity throughout the video. To generate this dataset, we collected approximately 1-minute-long video clips from both our AER lab drive and various publicly available datasets. These videos were converted to a maximum frame rate of 30 FPS. Subsequently, we divided them into two subgroups: single-animal and multi-animal. We annotated each frame of each video using our SAM tracking annotation strategy (see below).

Our single-animal dataset, used for evaluating single-object tracking, encompassed seven diverse experimental setups all of which, besides one, were distributed through OpenBehavior Video Repository (edspace.american.edu/openbehavior/video-repository/video-repository-2/). The dataset included the following videos: 1) ‘Olfactory search chamber’ (donated by Matt Smear); 2) ‘Open field, grayscale’ (donated by Sam Golden); 3) ‘Open field, RGB’ (donated by Sam Golden); 4) ‘Simple chamber, red light’ (from the AER lab); 5) ’Elevated plus maze’ (donated by Zachary Pennington and Denise Cai); 6) ‘Operant chamber, mouse’ (donated by Zachary Pennington and Denise Cai); and 7) ‘Operant chamber, rat,’ acquired with a fisheye lens (donated by Michael McDannald).

Our multi-animal dataset for evaluating multi-object tracking encompassed five diverse experimental setups: 1) ’Operant chamber, mixed’ (donated by Sam Golden acquired via OpenBehavior); 2) ’Home cage, mixed grayscale’ (15); 3) ’Home cage, mixed RGB’ (donated by Sam Golden acquired via OpenBehavior); 4) ’Enriched cage, mixed infrared’ acquired with an infrared camera (from the AER lab); and 5) ’Large cage, white triplet’ (from the AER lab).

### Segment Anything Model (SAM)-guided annotation strategy

#### Image annotation

To annotate object masks both efficiently and cost-effectively, we leveraged the Segment Anything Model (SAM), developed by Meta (11), as a guide for mask generation. SAM–a deep neural network–is designed for interactive instance segmentation in computer vision. It is adept at converting simple user inputs into high-quality object masks in images.

To annotate our detection data, we developed a graphical user interface (GUI). The interface allows users to interact with images by specifying foreground/background points or bounding boxes. SAM then converts these points into precise instance masks. Our annotation process utilizes two of SAM’s prompting strategies: (1) Point prompts, where the user specifies a set of points to indicate an object’s foreground or background. (2) Bounding box prompts, where SAM is provided the object of interest with a bounding box, which are used for annotating tracking data efficiently.

The annotation tool input is a folder containing images, while its output is a Common Objects in Context (COCO)-style metadata file (23) with instance segmentation annotations for the images. The pipeline for annotating a single image is as follows: 1) the user specifies a foreground/background point using the right/left mouse click, 2) SAM converts the point prompt into an instance mask, 3) if the predicted mask is accurate, the user can press <SPACE> to proceed to the next animal in the image, or <ESC> to move to the next image. If the mask is incorrect, the user can return to step 1 and refine the prompt, prompting SAM to update the mask based on the latest set of points.

### Tracking data annotation

Annotating tracking data poses significant time and cost challenges due to the large number of frames requiring annotation in each video (e.g., a 1-minute video at 25 FPS results in 1,500 frames). To expedite this process, we annotate frames sequentially while initializing the annotations for a current frame by providing SAM with the previous frame’s mouse bounding boxes and the current frame’s image. This method bootstraps annotation by taking advantage of the minimal movement of mice between frames, requiring only minimal further adjustments to the bounding boxes.

### Evaluation Procedures

#### Zero-shot evaluation

This strategy aims to assess the effectiveness of a model on a new, downstream task without any fine-tuning specific to that task. In this study, we begin all zero-shot analysis with a pretrained DAMM detector and directly evaluate its performance on the evaluation set.

#### Few-shot evaluation

This strategy aims to assess a model’s effectiveness on a downstream task when it has been exposed to a limited number of examples from that task. In this study, we conducted few-shot analyses with N ranging from 5 to 50 across various experiments. In these cases, we used the N examples to fine-tune the DAMM detector before its evaluation on the downstream task.

### Evaluation Metrics for detection

#### Mask Average Precision (AP) 75

Mask AP 75 in detection tasks evaluates the accuracy of instance segmentation, specifically measuring how well the model identifies instances of objects within an image by comparing the predicted masks to the ground truth masks. A mask is considered correctly identified if there is greater than 75% overlap with the ground truth mask.

We use evaluation metrics implemented in Detectron2.

### Evaluation Metrics

#### Single-object tracking accuracy

Single-object tracking accuracy (TA) assesses how accurately a model tracks a single object in video sequences. It is calculated using the following equation: TA = number of correctly tracked frames / total number of frames. For this paper we consider an IoU greater than 0.5 to be considered correctly tracked.

#### Multi object tracking accuracy

Multi-object tracking accuracy (MOTA) assesses how accurately a model tracks multiple objects in video sequences. The primary distinction from single object tracking accuracy is the inclusion of ID switches in the assessment. The calculation is as follows: MOTA = 1 - ((false negatives + false positives + id switches) /ground truth). For this paper we consider an IoU greater than 0.5 to be considered correctly tracked.

## Acknowledgments

We thank the members of the Johnson and the Eban-Rothschild labs for valuable discussion and feedback. We thank the members of the AER lab for collecting and sharing video data. This study was supported by the MICDE Catalyst Grant (to A.E.-R. and J.J.) and the National Institute of Neurological Disorders and Stroke (R01NS131821 and R01NS129874 to A.E.-R.).

## Author contributions

G.K., J.J. and A.E.-R. conceived and designed the study. G.K. and J.M. performed research. G.K. and A.E.-R. wrote the manuscript, with feedback from authors. J.J. and A.E.-R. supervised the study.

## Data, Materials, and Software Availability

The publicly available datasets are detailed in the Methods. DAMM weights will be available at publication time at github.com/backprop64/DAMM.

## References

1. A. Mathis, S. Schneider, J. Lauer, M. W. Mathis, A Primer on Motion Capture with Deep Learning: Principles, Pitfalls, and Perspectives. Neuron 108, 44–65 (2020).

2. T. D. Pereira, J. W. Shaevitz, M. Murthy, Quantifying behavior to understand the brain. Nat Neurosci 23, 1537–1549 (2020).

3. A. I. Dell et al., Automated image-based tracking and its application in ecology. Trends Ecol Evol 29, 417–428 (2014).

4. J. Lauer et al., Multi-animal pose estimation, identification and tracking with DeepLabCut. Nat Methods 19, 496–504 (2022).

5. T. D. Pereira et al., SLEAP: A deep learning system for multi-animal pose tracking. Nat Methods 19, 486–495 (2022).

6. J. M. Graving et al., DeepPoseKit, a software toolkit for fast and robust animal pose estimation using deep learning. eLife 8, e47994 (2019).

7. J. J. Sun et al., Self-Supervised Keypoint Discovery in Behavioral Videos. Proc IEEE Comput Soc Conf Comput Vis Pattern Recognit 2022, 2161–2170 (2022).

8. C. Weinreb, et al., Keypoint-MoSeq: parsing behavior by linking point tracking to pose dynamics. *bioRxiv* 10.1101/2023.03.16.532307 (2023).

9. M. I. Sotelo et al., Neurophysiological and behavioral synchronization in group-living and sleeping mice. Curr Biol 10.1016/j.cub.2023.11.065 (2023).

10. M. I. Sotelo et al., Lateral hypothalamic neuronal ensembles regulate pre-sleep nest- building behavior. Curr Biol 32, 806–822.e807 (2022).

11. A. Kirillov, et al., Segment anything. arXiv preprint arXiv:2304.02643 (2023).

12. A. Gupta, P. Dollar, R. Girshick (2019) Lvis: A dataset for large vocabulary instance segmentation. in *Proceedings of the IEEE/CVF conference on computer vision and pattern recognition*, pp 5356–5364.

13. K. He, G. Gkioxari, P. Dollár, R. Girshick (2017) Mask r-cnn. in *Proceedings of the IEEE international conference on computer vision*, pp 2961–2969.

14. A. Bewley, Z. Ge, L. Ott, F. Ramos, B. Upcroft (2016) Simple online and realtime tracking. in 2016 *IEEE international conference on image processing (ICIP)* (IEEE), pp 3464-3468.

15. J. J. Sun et al., Caltech Mouse Social Interactions (CalMS21) Dataset (1.0) [Data set]. CaltechDATA 10.22002/D1.1991 (2021).

16. S. Ye, et al., SuperAnimal models pretrained for plug-and-play analysis of animal behavior. arXiv preprint arXiv:2203.07436 (2022).

17. S. A. Golden, H. E. Covington, 3rd, O. Berton, S. J. Russo, A standardized protocol for repeated social defeat stress in mice. Nat Protoc 6, 1183–1191 (2011).

18. R. E. Kalman, A new approach to linear filtering and prediction problems. (1960).

19. H. W. Kuhn, The Hungarian method for the assignment problem. Naval Research Logistics Quarterly 2, 83–97 (1955).

20. Y. Wu, A. Kirillov, F. Massa, W.-Y. Lo, R. Girshick, Detectron2. (2019).

21. M. Abadi, et al., Tensorflow: Large-scale machine learning on heterogeneous distributed systems. *arXiv preprint arXiv:1603.04467* (2016).

22. J. Lauer et al., maDLC Tri-Mouse Benchmark Dataset - Training [Data set]. . *Zenodo*. 10.5281/zenodo.5851157 (2022).

23. T.-Y. Lin et al. (2014) Microsoft coco: Common objects in context. in *Computer Vision– ECCV 2014: 13th European Conference, Zurich, Switzerland, September 6-12*, *2014, Proceedings, Part V 13* (Springer), pp 740-755.

