## Supplementary data for "DAMM for the detection and tracking of multiple animals within complex social and environmental settings"

#### Supplementary Material

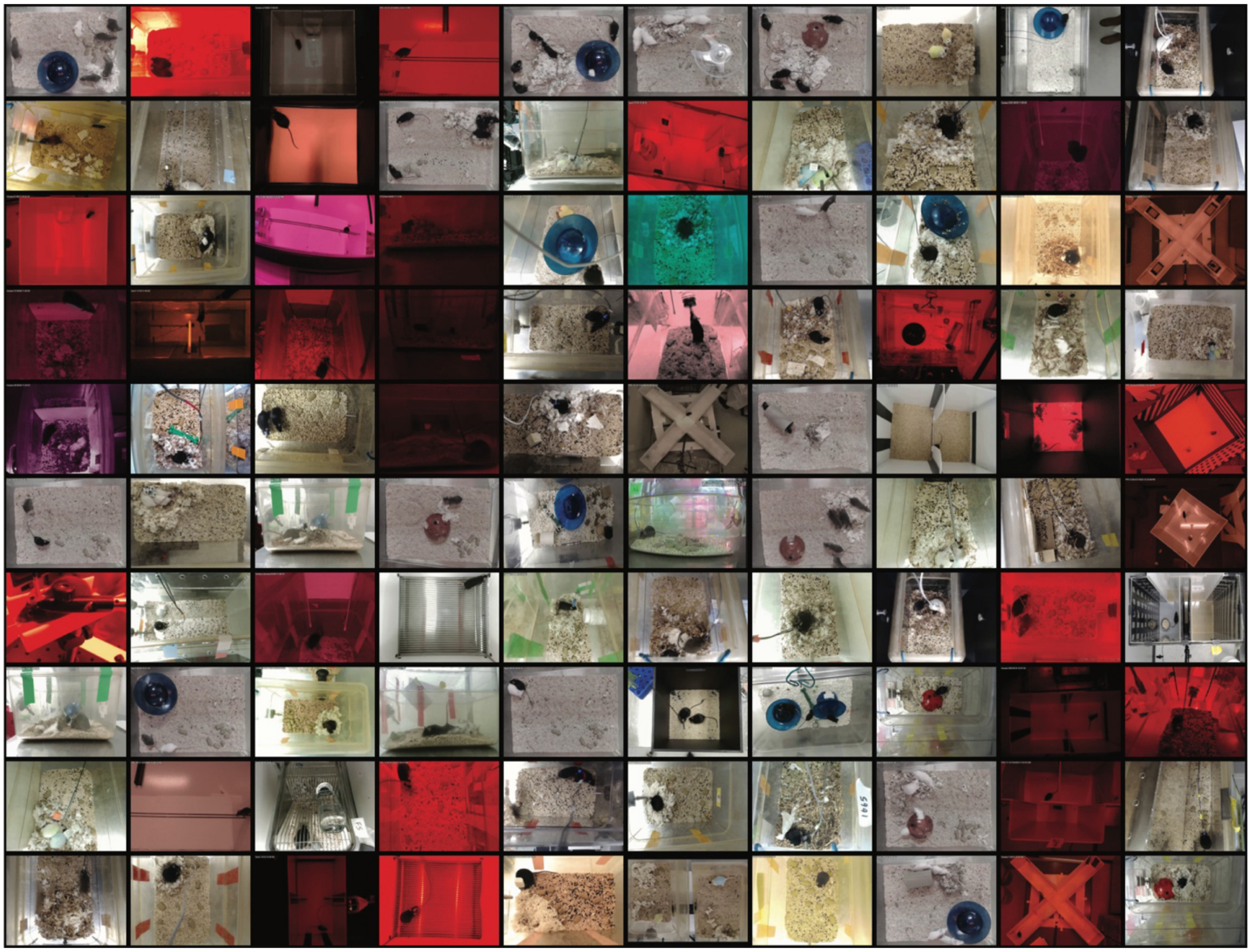

**Fig. S1:** A 10 x 10 grid of images from the AER-LG dataset showcasing diversity in terms of viewing angles, number of animals, coat colors, setup architectures and lighting conditions.

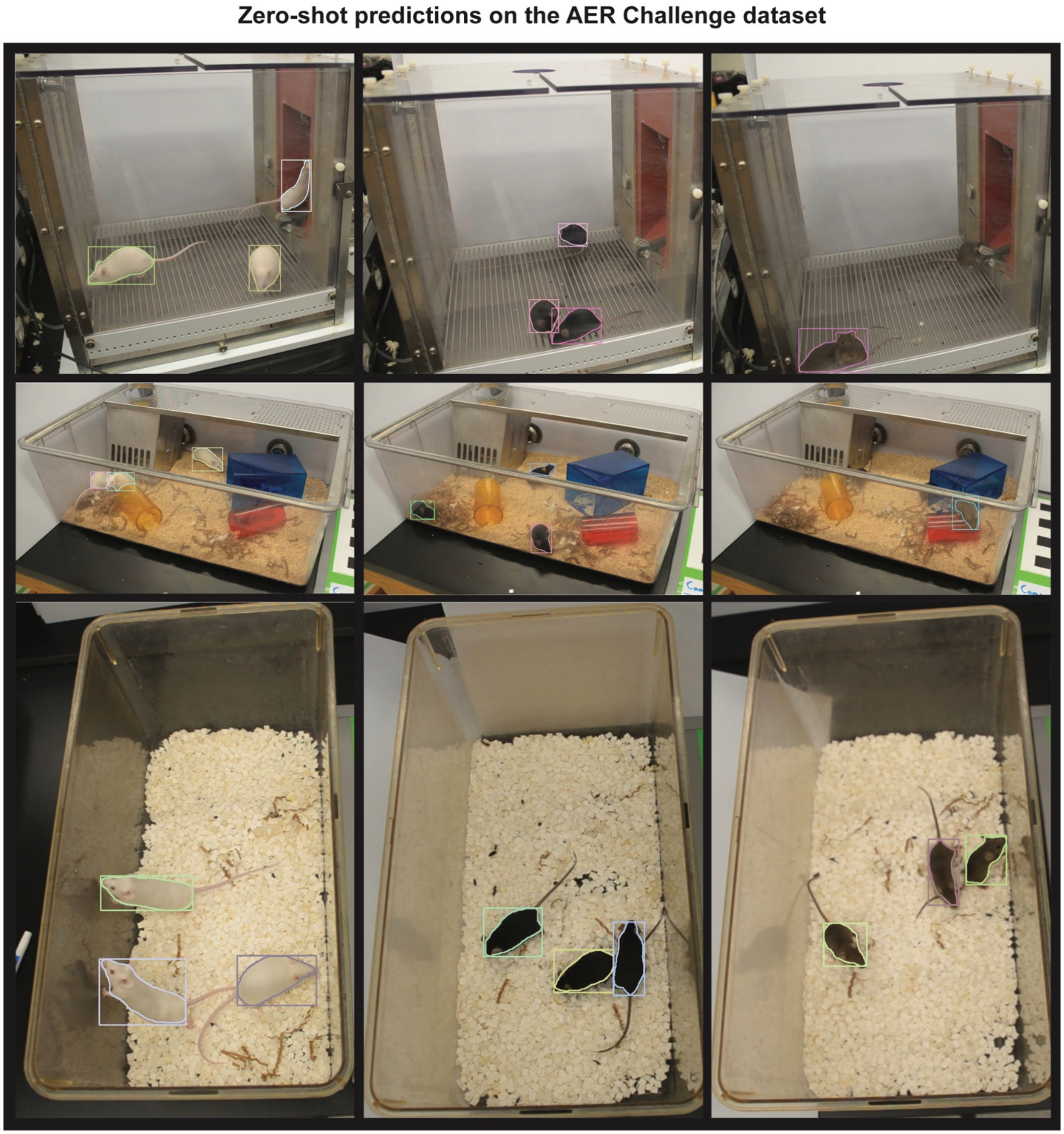


**Fig. S2:** A 3 x 3 grid displaying images from the high-end camera subset from the AER Challenge dataset, illustrating three types of enclosure architectures in separate rows (top: 'Large Cage,' middle: 'Operant Chamber,' and bottom: 'Enriched Cage'), and the different coat colors (first: white, second: black, and third: agouti).

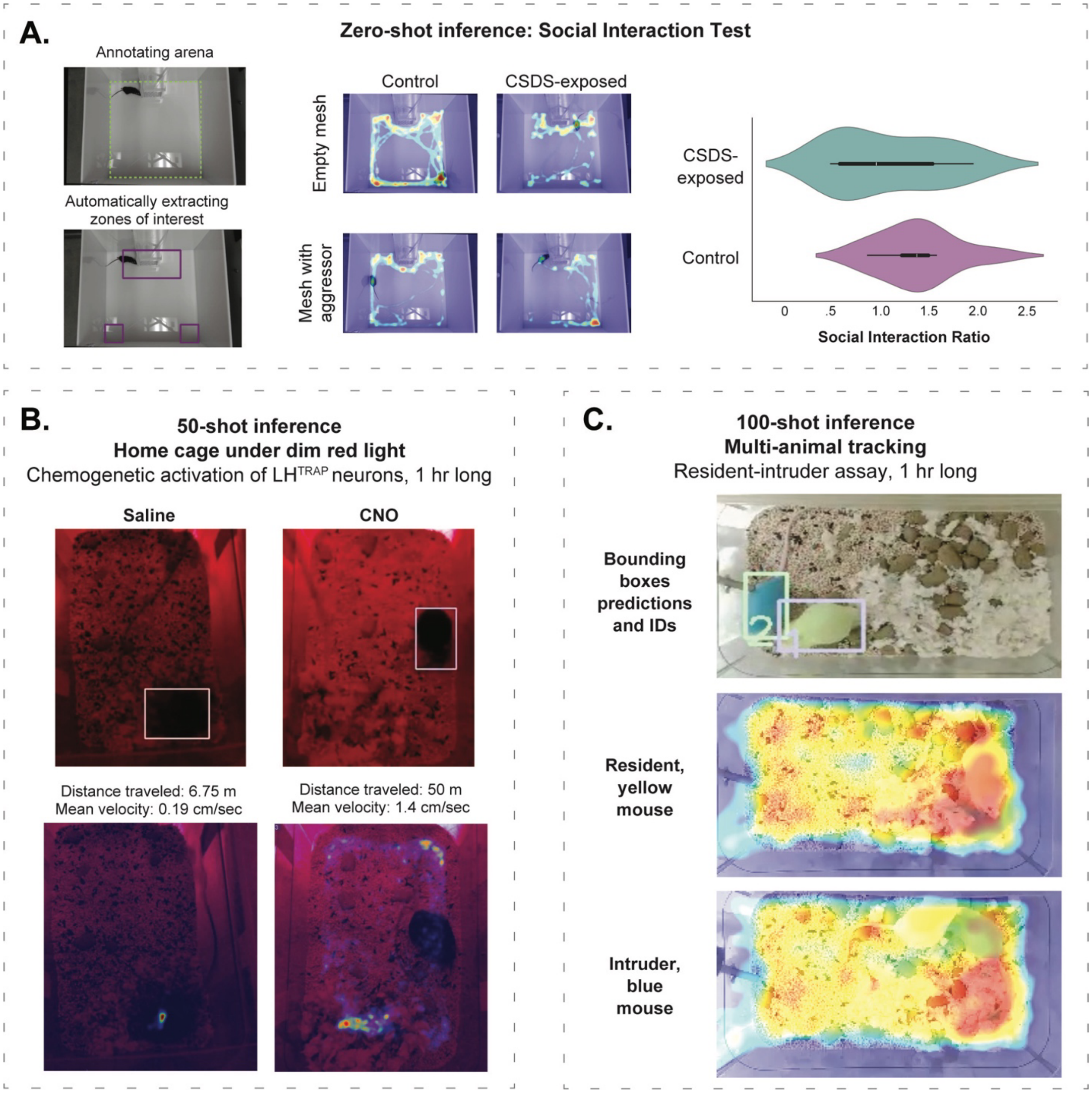


**Fig. S3: Application of DAMM in behavioral assays.**
(**A**) Application of DAMM zero-shot inference for the analysis of SI test data, for chronic social defeat stress (CSDS) exposed mice (n = 18 male C57 mice) and unstressed, control mice (n = 9 male C57 mice). Left, frames of the arena annotated using a Colab GUI, along with visualizations of the interaction and corner zones extracted automatically. Middle, Representative heatmaps of mouse location throughout the video during the SI test for an individual unstressed, control mouse (left) and a stressed, test mouse (right), prior (top) and following (bottom) the placement of a CD-1 aggressor in the mesh enclosure. Right, Violin plot illustrating the social interaction ratio in control and test mice. The plot's tubular bubble shape represents the data points' density distribution. Notably, the CSDS-exposed mice exhibit a bimodal distribution, indicating the presence of known susceptible and resilient phenotypes. The

median is marked by a white dot. The bold bar's lower end signifies the data's first quartile, and its upper end represents the third quartile. (**B**) Use of DAMM 50-shot inference for locomotion data analysis post-chemogenetic activation of pre-sleep $LH^{TRAP}$ neurons under dim-red light. The top section shows a random frame from DAMM tracking visualization post-administration of Saline (control, left) and CNO (test, right). The bottom section presents heatmaps of each mouse's location throughout a 1-hour period after drug administration. (**C**) Use of DAMM 100-shot inference for multi-animal tracking within a resident-intruder assay. The top section shows a random frame from DAMM tracking visualization showcasing bounding boxes and corresponding ID's. The bottom section presents heatmaps of each mouse's location throughout a 1-hour period after the placement of the intruder mouse into the home cage of the resident mouse (represented as log-transformed histogram counts). We manually corrected seven ID switched throughout the video.

### Supplementary Materials and Methods

#### Behavioral experiments details

##### Chronic Social Defeat Stress

###### *Mice*

We utilized male black C57BL/6J mice aged 8-12 weeks (bred in-house) and 6-8 month old ex-breeder male white CD-1 mice (Strain #: 022, sourced from Charles River Laboratories). The mice were housed in a controlled environment, maintained at a temperature of 22 ± 1°C with a 12-hour light/dark cycle and ad libitum access to food and water. Prior to the experiment, the mice were also provided with nesting material. All experiments were conducted in accordance with the US National Institutes of Health Guide for the Care and Use of Laboratory Animals and approved by the University of Michigan's Institutional Animal Care and Use Committee.

###### *Chronic Social Defeat Stress procedure*

We implemented a chronic social defeat stress (CSDS) model as described in Golden et al., 2011, in adult male C57BL/6J mice to induce stress-related phenotypes. The CSDS procedure spanned 10 consecutive days. Each day, a test mouse was introduced into the home cage (50 cm x 25 cm x 40 cm) of a novel aggressive CD-1 mouse for a period of 5-10 minutes, ensuring direct but controlled aggressive interactions. Post confrontation, the test mouse was separated from the aggressor by a transparent, perforated divider within the same cage, allowing visual, olfactory, and auditory contact for the remaining 24 hours (Golden et al., 2011). The aggressor mice were selected based on their established history of aggressive behavior, screened prior to the experiment. Control mice (adult male C57BL/6J of a similar age) were left undisturbed in their home cages for 10 days. CSDS-exposed and control mice were transferred to new cages on day 11.

###### *Social interaction (SI) test*

Approximately 23 hours after the transfer of the CSDS-exposed and control mice to new cages, they were subjected to a SI test to evaluate the behavioral impacts of chronic social defeat (Golden et al., 2011). This test aimed to assess changes in social behavior potentially induced by the CSDS experience. The test was conducted in an arena measuring 44.5 cm by 44.5 cm, divided into two consecutive 150-second phases. In the first phase, the test mouse was introduced into the arena containing an empty wire mesh cage (10 cm by 6.5 cm), allowing for baseline sociability observations. Subsequently, the test mouse was gently removed, and an unfamiliar CD-1 mouse was placed inside the wire mesh cage. In the second phase, the test mouse was reintroduced to the arena, now with the CD-1 mouse present in the cage, to assess changes in social behavior. All trials were video recorded using high-definition webcams (either Logitech C920 or Angetube 1080p), positioned above the arena. To calculate the SI ratio, the time a mouse spent in the interaction zone with a target CD-1 present was divided by the time it spent in the interaction zone when a target CD-1 was absent.

#### Chemogenetic activation of LH-TRAPed neurons

##### *Mice*

We utilized reproductively inexperienced F1 $Fos^{2A\text{-}iCreERT2}$ (TRAP2; The Jackson Laboratory, Stock #: 030323) mice > 8 weeks old weeks (bred in-house by crossing with black C57BL/6J mice). The mice were housed in a controlled environment, maintained at a temperature of 22 ± 1°C with a 12-hour light/dark cycle and ad libitum access to food and water. The mice were provided with compressed cotton 'Nestlet' nesting material (Ancare, Bellmore, NY, U.S.A.), shredded paper 'Enviro-Dri' nesting material (Shepherd Specialty Papers, Watertown, TN, U.S.A.). During the experiment, mice were individually housed in custom Plexiglas recording chambers (28.6 × 39.4 cm and 19.3 cm high). All experiments were conducted in accordance with the US National Institutes of Health Guide for the Care and Use of Laboratory Animals and approved by the University of Michigan's Institutional Animal Care and Use Committee.

Mice were anesthetized with a ketamine-xylazine mixture (100 and 10 mg kg-1, respectively; intraperitoneal injection, IP) and administered with lidocaine and carprofen (4 mg kg-1 and 5 mg kg-1, respectively). Mice were placed into a stereotaxic frame (David Kopf Instruments, Tujunga, CA, U.S.A.) and maintained under isoflurane anesthesia (~1% in O2). We stereotaxically infused viral vectors (AAV-EF1α-DIO-hM4Gq-mCherry) into the lateral hypothalamus (AP = -1 mm, ML = ±1.15 mm and DV = −4.9 mm) at a slow rate (100 nl min-1) using a microinjection syringe pump (UMP3T-1, World Precision Instruments, Ltd.) and a 33G needle (Nanofil syringe, World Precision Instruments, Ltd.). After infusion, the needle was kept at the injection site for ≥ 8 min and then slowly withdrawn. The skin was then closed with surgical sutures. Mice were placed on a heating pad until fully mobile. Following recovery from surgery (~10 days), mice were separated into individual recording chambers.

Mice were acclimated to handling and intraperitoneal (IP) injections for approximately one week prior to the experiment. On the day of the experiment, starting at Zeitgeber Time (ZT) 0, the nests in the home cages of the test mice were dispersed. This was followed by 4-hydroxytamoxifen (4-OHT) administration at ZT 1. Subsequently, the original nests were removed, and the mice were provided with fresh nesting material. This intervention extended their pre-sleep phase. Throughout the 2-hour period following the 4-OHT administration, an experimenter monitored the mice continuously to prevent them from sleeping, supplying additional nesting material as needed to keep them engaged and awake. After this period, from ZT 2 to 24, the mice were left undisturbed.

##### *Chemogenetic manipulation*

Mice were removed from their home cages at the beginning of the dark phase (ZT 12), IP administered either saline or CNO (1 mg kg-1), and returned to the home cage with as little disturbance to the nest as possible. Mice behavior was video recorded using high-definition webcams (either Logitech C920 or Angetube 1080p).

#### Resident-intruder test

##### *Mice*

We utilized male white CD-1 mice (bred in-house). The mice were housed in a controlled environment, maintained at a temperature of 22 ± 1°C with a 12-hour light/dark cycle and ad libitum access to food and water. The fur of the mice was dyed with either Blue Moon (blue) or Electric Lizard (green) dyes from Tish & Snooky's Manic Panic (manicpanic.com). All experiments were conducted in accordance with the US National Institutes of Health Guide for the Care and Use of Laboratory Animals and approved by the University of Michigan’s Institutional Animal Care and Use Committee.

##### *Experimental procedure*

Mice were individually housed for approximately one week prior to the experiment. At ZT 0, a male non-sibling "intruder" was placed into the home-cage of a "resident" mouse. Mice behavior was video recorded using high-definition webcams (either Logitech C920 or Angetube 1080p).
